# Phototropin-mediated perception of light direction in Arabidopsis leaves regulates blade flattening

**DOI:** 10.1101/2021.05.25.445665

**Authors:** Martina Legris, Bogna Maria Szarzynska-Erden, Martine Trevisan, Laure Allenbach-Petrolati, Christian Fankhauser

## Abstract

One conserved feature among angiosperms is the development of flat thin leaves. This developmental pattern optimizes light capture and gas exchange for photosynthesis. The blue light receptors phototropins are required for leaf flattening, with the null *phot1phot2* mutant showing downwards curled leaves in Arabidopsis. However, key aspects of their function in leaf development remain unknown. Here, we performed a detailed spatiotemporal characterization of phototropin function in Arabidopsis leaves. We found that phototropins perceive light direction in the leaf blade, and similar to their role in hypocotyls they control the spatial pattern of auxin signaling possibly modulating auxin transport, to ultimately regulate cell expansion. Phototropin signaling components in the leaf partially differ from hypocotyls. Moreover, the light response on the upper (adaxial) and lower (abaxial) sides of the leaf blade suggest a partially distinct requirement of phototropin signaling components on each side. In particular, NON PHOTOTROPIC HYPOCOTYL 3 (NPH3) showed an adaxial-specific function. In addition, we show a prominent role of PHYTOCHROME KINASE SUBSTRATE 3 (PKS3) in leaf flattening. Among the auxin transporters tested, PINs and AUX/LAX influence the response most prominently. Overall, our results show that directional blue light perception by the phototropins is a key aspect of leaf development, integrating endogenous and exogenous signals.

**One sentence summary:** Phototropins perceive light direction in the leaf and control the auxin signaling pattern to regulate blade flattening.

## INTRODUCTION

Plants are photoautotrophic organisms, and as such optimization of light capture is key for their success. In most land plants leaves are the major photosynthetic organ (Chitwood and Sinha, 2016). Leaves start developing in the shoot apical meristem, where following an auxin response maxima a primordium differentiates. Primordia are patterned early with different gene expression domains in the portion closer to the meristem (adaxial side) versus distally from the meristem (abaxial side) and in the medio-lateral and proximo-distal axes. In the juxtaposition between the adaxial and abaxial side, the marginal domain will promote leaf expansion. Early in development cells divide, and in Arabidopsis an arrest front in cell division moves from the tip to the base of the leaf. Subsequently, cells expand and a growth arrest front follows the same pattern from tip to base (Xiong and Jiao, 2019; Heisler and Byrne, 2020). In summary, leaf shape depends on the early patterning of the primordia, cell division rates and coordinated expansion of cell layers.

One conserved feature of angiosperm leaves is their flat shape, with wide thin blades. This shape optimizes light capture, as well as gas exchange and temperature regulation (Inoue et al., 2008; de Carbonnel et al., 2010). Thus, it is not surprising that in addition to endogenous cues, environmental signals, in particular light, also control leaf flattening. The red (RL) and far-red light receptor phytochrome B (phyB) promotes leaf curling, and in shade conditions inactivation of the receptor triggers blade flattening (Kozuka et al., 2012; Johansson and Hughes, 2014). Blue light (BL) perceived by phototropins (phot1 and phot2 in Arabidopsis) promotes leaf expansion and flattening (Sakai et al., 2001; Takemiya et al., 2005). However, whether phototropins act as sensor of light direction to control blade flattening is currently not known.

Phototropins are membrane-associated serine-threonine kinases containing two light, oxygen and voltage (LOV) domains that allow them to perceive BL and ultraviolet light (Legris and Boccaccini, 2020). Their role in stem phototropism has been studied in depth. When the light stimulus is directional, a light gradient is established within the stem creating a gradient of phototropin activation. Activated phototropins are autophosphorylated and at the plasma membrane interact with members of the NPH3/RPT2-Like (NRL) and Phytochrome Kinase Substrate (PKS) families. NPH3 and RPT2 have a role in stem phototropism, and mutants in these genes also have a defect in leaf flattening (Inoue et al., 2008; de Carbonnel et al., 2010; Harada et al., 2012; Christie et al., 2018). It has been shown that NPH3 is phosphorylated by phot1 as an early step of the signaling pathway (Sullivan et al., 2021). Another direct phototropin phosphorylation target is PKS4 (Demarsy et al., 2012). Among PKS proteins, PKS1 and PKS4 have a role in stem phototropism, while PKS2 has a role in leaf flattening (de Carbonnel et al., 2010; Kami et al., 2014). While the molecular function of these proteins remains unknown, they have been related to auxin transport or signaling (de Carbonnel et al., 2010; Kami et al., 2014). In the stem, a few minutes after unilateral irradiation an auxin gradient is created throughout the organ. Higher auxin concentrations on the shaded side of the stem promote cell expansion, while growth on the lit side is inhibited causing bending towards the light (Legris and Boccaccini, 2020). Auxin transporters of the PIN-formed (PIN) family are required for hypocotyl bending (Willige et al., 2013). In particular, in response to unilateral BL, PIN3 in the endodermis re-localizes to the outer membrane, allowing auxin transport from the vasculature to the epidermis, where organ growth is controlled (Ding et al., 2011). Another auxin transporter, ABCB19, is a phosphorylation target of phot1, and regulates hypocotyl phototropism by promoting basipetal auxin fluxes (Christie et al., 2011).

The plant hormone auxin has a strong role during leaf development (Xiong and Jiao, 2019; Heisler and Byrne, 2020). As mentioned before, leaves start developing in sites with high auxin signaling in the meristem. Recently a role for auxin has been proposed in the establishment and/or maintenance of the adaxial-abaxial polarity. Lower auxin in the adaxial side of the blade is proposed to promote the adaxial fate, allowing flattening (Guan et al., 2017). Auxin signaling in the middle domain is required for leaf expansion. In the adult leaf auxin is synthesized in the margins and transported towards the stem. In accordance with these roles, several mutants in auxin synthesis, signaling or transport genes fail to grow flat leaves, and exogenous application of auxin or auxin signaling inhibitors promote rolling of the blade (Watahiki and Yamamoto, 1997; de Carbonnel et al., 2010; Jenness et al., 2019; Xiong and Jiao, 2019; Heisler and Byrne, 2020; Jenness et al., 2020).

Provided that leaf development relies on internal cues, auxin being one of them, and that phototropins regulate leaf shape and can regulate auxin signaling, we set out to evaluate the mechanisms underlying the interaction between light perception and the leaf developmental program to regulate blade flattening. Using specific light treatments, we performed a detailed spatiotemporal analysis of the effect of phototropin activation on leaf shape. We show that throughout leaf development, in addition to internal cues, light direction perceived by phototropins regulates cell expansion to regulate blade morphology.

## RESULTS

### Leaf shape depends not only on the perception of BL, but also on its site of perception

BL perceived by phototropins is required to grow flat leaves (Takemiya et al., 2005; de Carbonnel et al., 2010; Kozuka et al., 2012; Jenness et al., 2020). However, whether this depends on phototropins’ ability to sense light direction remains unclear. In most plants, but in particular in Arabidopsis which is a rosette, leaves intercept light with the adaxial side, and most of it is absorbed by the photosynthetic pigments, creating a steep blue light gradient throughout the leaf (Vogelmann et al., 1989; Paradiso et al., 2020). Given that a similar signal perceived by phototropins triggers phototropism in the stem, we evaluated whether phototropins also perceive light direction in the leaf. We measured the leaf flattening index (LFI), as the ratio between the leaf blade area before and after artificially flattening it (de Carbonnel et al., 2010), in plants grown in RL with addition of 0,1 µmol.m^−2^.s^−1^ of BL either from the top or from below (Fig. 1A, B). While plants (WL) grown with 0,1 µmol.m^−2^.s^−1^ BL from the top grew flat leaves (Fig. 1A), those grown in RL plus BL from below failed to grow flat leaves and had an even stronger flattening defect than those grown in RL in the absence of BL (Fig. 1B). Since plants grown on plates had a different morphology than soil-grown plants (Fig. S1B), we also performed similar analyses in the latter, more natural, conditions. We covered the soil surface with colored aluminum foil, which reflects ambient light, irradiating the abaxial side of leaves (Fig. S1C). Dark aluminum foil did not affect leaf development compared to uncovered soil (Fig. S1A, B). Reflection of WL triggered downward bending of the leaf blade (Fig. 1C, G, S1C). As observed on plates, reflection of BL was enough to interfere with normal leaf flattening, while reflection of RL or yellow light (YL) did not affect flattening (Fig. 1C, Fig. S1C). These results confirm that BL is required to grow flat leaves and show that the site of BL perception in the leaf affects its shape.

**Figure 1 –.**
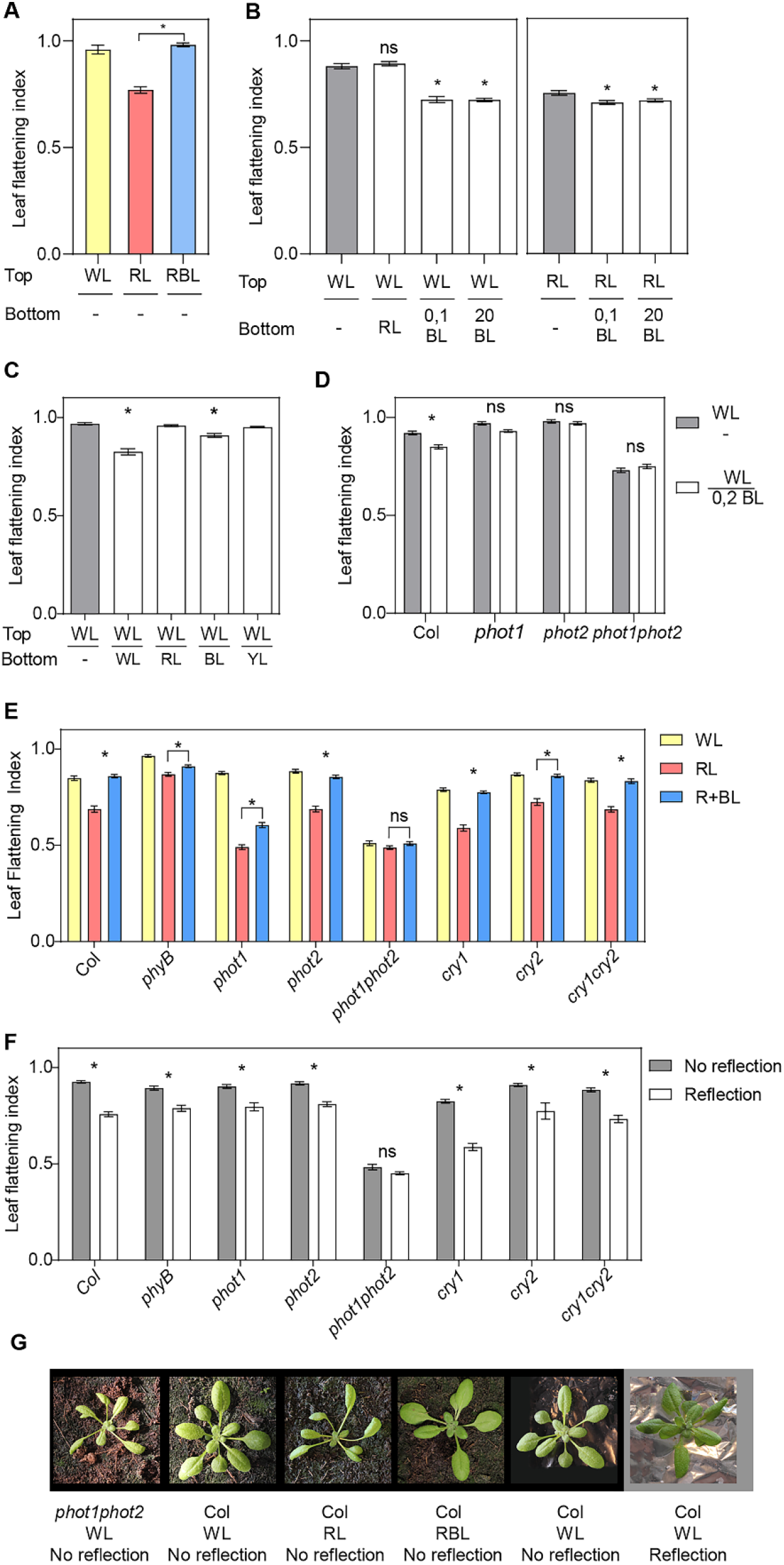
Leaf shape responds to light direction in a phototropin-dependent manner. A- Leaf flattening index in Col plants grown in plates for three weeks in white light (WL), red light (RL) or red + blue light (RBL). Bars represent mean ±SE of 16 - 30 leaves. * p < 0.05 in an unpaired T test. B- Leaf flattening index in Col plants grown in plates for two weeks in white light and transferred to WL or RL from the top supplemented with RL, 0,1 µmol.m^−2^.s^−1^ blue light (BL) or 20 μmol.m^−2^.s^−1^ BL from the bottom or dark bottom (-) for three days. Bars represent mean ±SE of 56 - 60 leaves. * p < 0.05 in ANOVA followed by Dunnett’s multiple comparison test. In each case, treatments involving light from below were compared to the same light from the top (WL or RL) without light from below (grey bars). C- Leaf flattening index in Col plants grown in WL for three weeks in soil covered with aluminum foil which reflects various colors (see spectra of the reflected light in Fig. S1B). YL: yellow light. Bars represent mean ±SE of 30 - 88 leaves. * p < 0.05 in ANOVA followed by Dunnett’s multiple comparison test. Treatments involving light from below were compared to the treatment without light from below (-, grey bar). D- Leaf flattening index in Col and phototropin mutants grown in plates in WL for two weeks and transferred to WL from the top and darkness (-) or 0,2 μmol.m^−2^.s^−1^ BL from the bottom for three days. Bars represent mean ±SE of 34 - 92 leaves. * p < 0.05 in ANOVA followed by Dunnett’s multiple comparison test between the two treatments for each genotype. E- Leaf flattening index in Col and photoreceptor mutants grown in soil for two weeks and transferred to WL, RL or RBL for one week. Bars represent mean ±SE of 36 - 50 leaves. * p < 0.05 in ANOVA followed by Dunnett’s multiple comparison test. F- Leaf flattening index in Col and photoreceptor mutant plants grown in WL for three weeks in soil covered with dark (No reflection) or clear (Reflection) aluminum foil. Bars represent mean ±SE of 16 – 29 plants. *p < 0.05 in ANOVA followed by Dunnett’s multiple comparison test. G- Representative pictures of *phot1phot2* and Col plants grown in selected light conditions.

### Phototropins perceive light direction in the leaf

In agreement with previous reports (Takemiya et al., 2005; de Carbonnel et al., 2010; Kozuka et al., 2012; Jenness et al., 2020), phototropins were the main photoreceptors perceiving BL and promoting flattening when BL reaches the adaxial side of the leaf (Fig. 1E, G). This was also the case when the abaxial side of the leaf was irradiated (Fig. 1F). When plants were grown on plates, both phot1 and phot2 were required to respond to very low BL intensities (Fig. 1D). When plants were grown on soil, both in response to light from the top or below phot1 and phot2 triggered responses to low BL intensities and only the double mutant *phot1phot2* showed no BL response (Fig. 1 E, F).

phyB influences flattening in an opposite way to the phototropins (Kozuka et al., 2012). In our conditions *phyB* mutants were not very different from Col in WL, although, consistent with previous findings, *phyB* leaves were flatter than Col particularly in RL (Fig. 1E) (Kozuka et al., 2012). However, phyB did not significantly affect the response to light from below, showing that this light receptor does not contribute to the perception of light direction (Fig. 1F). It was recently reported that some leaf development defects in *phyB-9* were caused by a second mutation in VENOSA4 present in the *phyB-9* allele (*ven4-bnen*) (Yoshida et al., 2018), but this was not the case for leaf flattening (Fig. S1D). Interestingly, cry1 had an effect on leaf flattening, with the *cry1-304* mutant showing slightly curled leaves when light was perceived on the adaxial side and responding more to light from below (Fig. 1E, F). However, the double *cry1cry2* mutant did not show this effect, and in all cases *cry* mutants responded to the light stimulus. Hence, cryptochromes may have a regulatory role but not a direct effect on the control of leaf shape by directional light. In summary, similar to what is known in hypocotyls and inflorescence stems, in the leaf blade phototropins perceive light direction and control blade curvature.

### The perception of light direction in the leaf involves differential phototropin activation in the adaxial and abaxial layers

In contrast to the radial symmetry of the hypocotyl, Arabidopsis leaves have a bilateral polarity with well-defined adaxial and abaxial domains (Xiong and Jiao, 2019; Heisler and Byrne, 2020). Taking advantage of this, we used site-specific promoters to express *PHOT1* on the adaxial or the abaxial side of the leaf. If leaf flattening is controlled by differential phototropin activation, changing the expression domain of phototropins is expected to influence the response to light direction. *phot1phot2* mutants were complemented with *PHOT1-citrine* expressed under the promoter of *ASYMMETRIC LEAVES 2* (*pAS2*) or *FILAMENTOUS FLOWER* (*pFIL*), inducing its expression predominantly in the adaxial or abaxial domains, respectively (Fig. S2B) (Guan et al., 2017; Xiong and Jiao, 2019). With light from the top most transgenic lines showed complementation of the *phot1phot2* flattening defect (Fig. 2A). However, lines expressing *PHOT1* on the adaxial side of the leaf showed better complementation, and among the three lines expressing *PHOT1* on the abaxial side of the leaf one complemented poorly and another one partially (*pFIL-9* and *pFIL-37* respectively). The site of *PHOT1* expression affected leaf flattening to light from the abaxial side more clearly. Lines expressing *PHOT1* predominantly in the adaxial side of the leaf (*pAS2* lines) did not respond to the abaxial light stimulus, while those expressing *PHOT1* predominantly in the abaxial side (*pFIL* lines) responded to the abaxial light stimulus more than *phot2* (Fig. 2B). Expressing *PHOT1* in the epidermal layers under the *MERISTEM LAYER 1* promoter (*pML1*) (Preuten et al., 2013) had a similar effect as expressing it in the whole blade under the *pPHOT1* promoter (Preuten et al., 2015), suggesting that, a phot1 activation gradient throughout the leaf blade was not required, but differential activation on each epidermal layer was sufficient to regulate blade shape. The response to light correlated with the *PHOT1* expression pattern and not only with *PHOT1* levels, since lines expressing similar levels showed contrasting responses (*pAS2-1* and *pFIL-31*) and lines expressing different levels showed the same trend (*pFIL-9* and *pFIL-37* or *pML1* and *pAS2-4*, Fig. S2A). However, we note that among the *pFIL* lines the expression level correlated with their ability to complement *phot1phot2* when light was coming from the top (Fig. 2A, S2A). Collectively, these results indicate that in the leaf a directional light stimulus leads to differential phototropin activation on the adaxial versus abaxial layers of the leaf which in turn control blade flattening.

**Figure 2 –.**
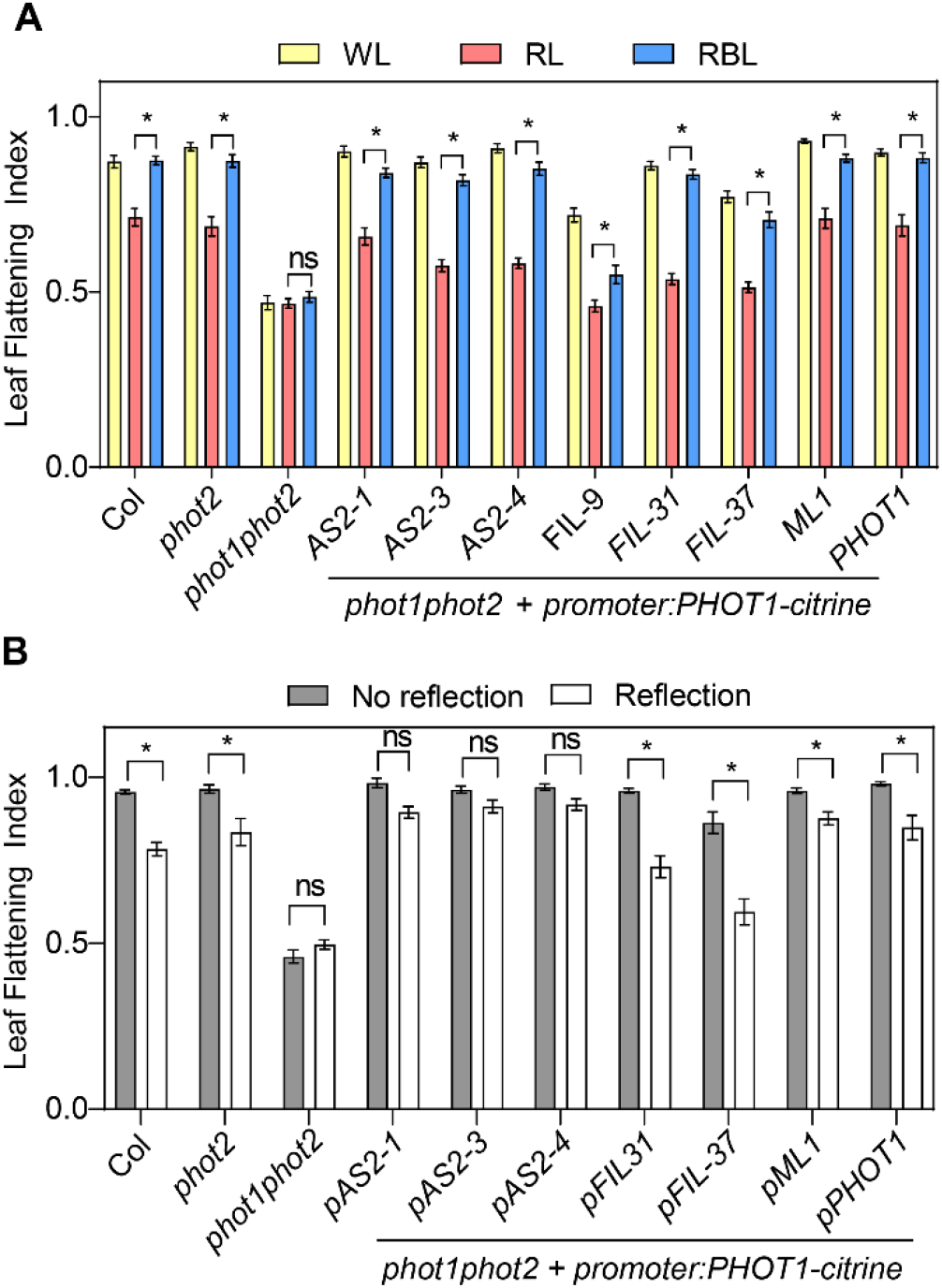
The PHOT1 expression domain regulates leaf shape. A- Leaf flattening index in plants grown in soil in WL for two weeks and transferred to WL, RL or RBL for one week. Bars represent mean ±SE of 17 - 28 leaves. * p < 0.05 in ANOVA followed by Dunnett’s multiple comparison test. B- Leaf flattening index in plants grown in WL in soil covered with dark (No reflection) or clear (Reflection) aluminum foil for three weeks. *pAS2-1*, *pAS2-3*, *pAS2-4*, *pFIL-31* and *pFIL-37* are independent transgenic lines expressing *PHOT1-citrine* under the promoters of *ASYMMETRIC LEAVES* (*AS*) or *FILAMENTOUS FLOWER* (*FIL*), respectively. *pAS2* drives *PHOT1* expression in the adaxial domain, and *pFIL* in the abaxial domain. Bars represent mean ±SE of 6 - 28 plants. * p < 0.05 in ANOVA followed by Dunnett’s multiple comparison test.

### Asymmetric requirement for some phototropin signaling elements in the control of leaf flattening

Early phototropin signaling during hypocotyl phototropism involves autophosphorylation of the receptor, in some cases followed by its degradation, and interaction with signaling members of the NRL and PKS families (Sakamoto and Briggs, 2002; Kong et al., 2006; Roberts et al., 2011; Christie et al., 2018; Legris and Boccaccini, 2020; Sullivan et al., 2021). We evaluated whether these mechanisms were conserved in response to BL in leaf blades. Consistent with observations in hypocotyls and in young leaves, in our conditions endogenous PHOT1 levels decreased in response to BL from the top, with a stronger effect of higher BL intensities (Fig. 3A, S3A) (Sakamoto and Briggs, 2002; Kozuka et al., 2011). A similar response was observed using a transgenic line expressing *pPHOT1:PHOT1-citrine* (Fig. S3B). In response to light irradiating the abaxial side a reduction in the fluorescent signal could be observed in the abaxial epidermis for *pML1:PHOT1-citrine*, but not for *pCER6:PHOT2-GFP* (Fig. 3B). This is consistent with higher stability of phot2 than phot1 in response to BL in hypocotyls and leaves (Sakamoto and Briggs, 2002; Kong et al., 2006; Kozuka et al., 2012).

**Figure 3 –.**
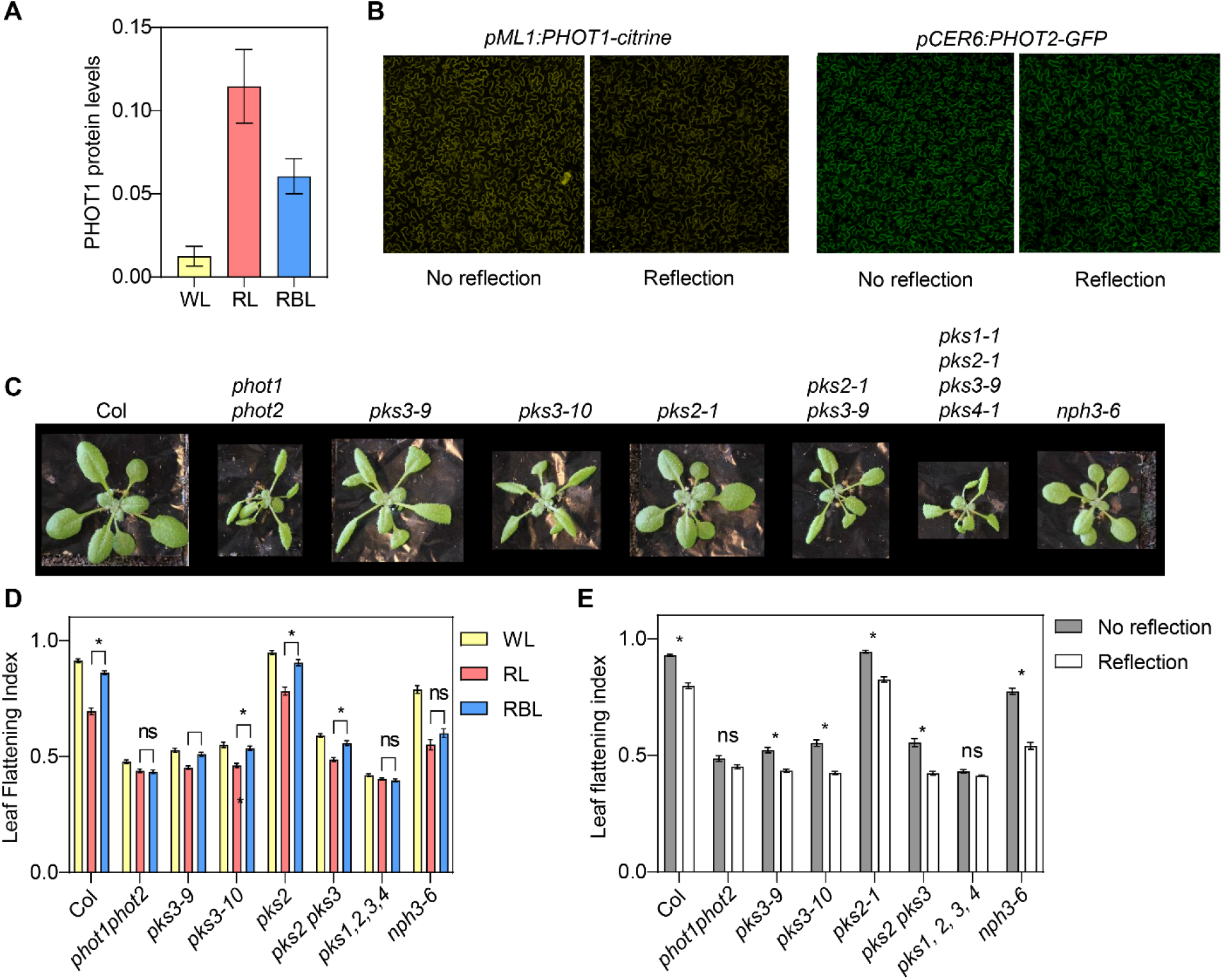
Early phototropin signaling mechanisms in leaves. A- PHOT1 levels in leaf blades of Col plants grown for two weeks in WL and transferred to WL, RL or RBL for one day. Protein blots were probed with anti PHOT1 antibody and DET3 was used as a loading control. Quantification of western blots shown in figure S3A. Each bar represents mean ±SE of four replicates. B- PHOT1 levels decrease in the abaxial epidermis in response to reflected light. Confocal images of plants expressing *pML1:PHOT1-citrine* or *pCER6:PHOT2-GFP* in the epidermis. Plants grown for three weeks in WL on soil covered with dark (No reflection) or clear (Reflection) aluminum foil. C- Pictures of Col and phototropin, *pks* and *nph3* mutant plants grown for three weeks in WL with no reflection. D- Leaf flattening index in plants grown on soil in WL for two weeks and transferred to WL, RL or RBL for one week. Bars represent mean ±SE of 22 - 56 leaves. * p < 0.05 in ANOVA followed by Dunnett’s multiple comparison test. E- Leaf flattening index in plants grown in WL on soil covered with dark (No reflection) or clear (Reflection) aluminum foil for three weeks. Bars represent mean ±SE of 20 - 45 leaves. * p < 0.05 in ANOVA followed by Dunnett’s multiple comparison test.

NRLs and PKS proteins regulate phototropin-mediated flattening in response to light from the top (Inoue et al., 2008; de Carbonnel et al., 2010; Harada et al., 2012). To compare the phototropin signaling network in the adaxial and the abaxial sides of the leaf, we measured LFI in response to light from the top and from below in *nph3* and *pks* mutants (Fig. 3C, D, E). *nph3* mutants showed partially impaired leaf flattening when grown in WL (Fig. 3C, D, E), but NPH3 was required to respond to low BL intensities coming from the top, as reported previously (Fig. 3D) (Inoue et al., 2008). However, when light was reflected and reached the abaxial side of the leaf, *nph3* mutants responded more than Col (Fig. 3E), suggesting that the early signaling mechanisms controlling leaf flattening in response to light differ between the adaxial and the abaxial sides of the blade. The PKS family has four members in Arabidopsis, but so far only *pks1*, *pks2* and *pks4* null mutants were available. To evaluate the role of *PKS3*, we used CRISPR to create two null alleles of PKS3, namely *pks3-9* and *pks3-10*. Interestingly, these mutants showed a striking flattening defect, comparable to *phot1phot2* mutants when grown in WL, suggesting that this member of the PKS family is a key phototropin signaling component in the leaf blade (Fig. 3C). However, both single mutant alleles showed a reduced but still significant response to light in all our conditions (Fig. 3D, E). *PKS2*, which was implicated in phototropin signaling in the leaf (de Carbonnel et al., 2010) did not show an effect in our conditions when mutated alone or in combination with *PKS3* (Fig. 3D, E). Nevertheless, the quadruple *pks1pks2pks3pks4* mutant showed an extreme flattening defect in all conditions tested, and did not respond to any light stimuli, showing that, while *PKS3* has a major role in leaf flattening, there is functional redundancy among PKS family members.

### Phototropins regulate leaf flattening reversibly and during the expansion phase

To change organ shape plants can regulate cell number, size or shape. During the first steps of leaf development the primordia undergoes cell division, followed by a front of cell expansion moving from the tip to the base of the blade (Xiong and Jiao, 2019). In order to determine whether phototropins control cell division or cell expansion in the leaf, we activated or inactivated phototropins at different times during development by transferring plants to WL, RL or RBL.

Plants grown in RL or RBL since day 0 after germination showed the same LFI as those transferred 6 or 9 days after germination, when leaf 1 and 2 are already more than 4mm long and cell division should be finished (Marrocco et al., 2009), suggesting that phototropins regulate leaf shape during the expansion phase (Fig. 4, S4). Consistent with this, LFI was comparable between Col and *phot1phot2* leaves during early development, and gradually *phot1phot2* curled downwards while Col leaves remained flat (Fig. 4B). In addition, the response to reflected light in Col plants occurred late in development and increased as leaves expanded (Fig. 4B). These results indicate that phototropins regulate leaf shape in large part by controlling cell expansion. To test this further, we used plants expressing the cell division marker *pCYCB1;1::NterCYCB1-GUS* (*cycB1;1-GUS*) (Marrocco et al., 2009) and stained them after 14 days of growth, prior to transfer to light treatments (Fig. 4C, D). We observed no GUS staining in leaves 1 and 2, which later responded to phototropin inactivation by a RL treatment, showing that leaf flattening was controlled during cell expansion (Fig. 4C, D). Finally, the response to light signals was reversible as long as the treatments occurred while the leaf was still expanding. Transferring plants from RL to RBL at day 12 or 15 after germination resulted in a higher LFI than keeping them in RL, but if the transfer was done at day 21 after germination BL could not promote leaf flattening (Fig. 4A, S4). We conclude that phototropin-mediated leaf flattening occurs at least in part through light-controlled cellular expansion.

**Figure 4 –.**
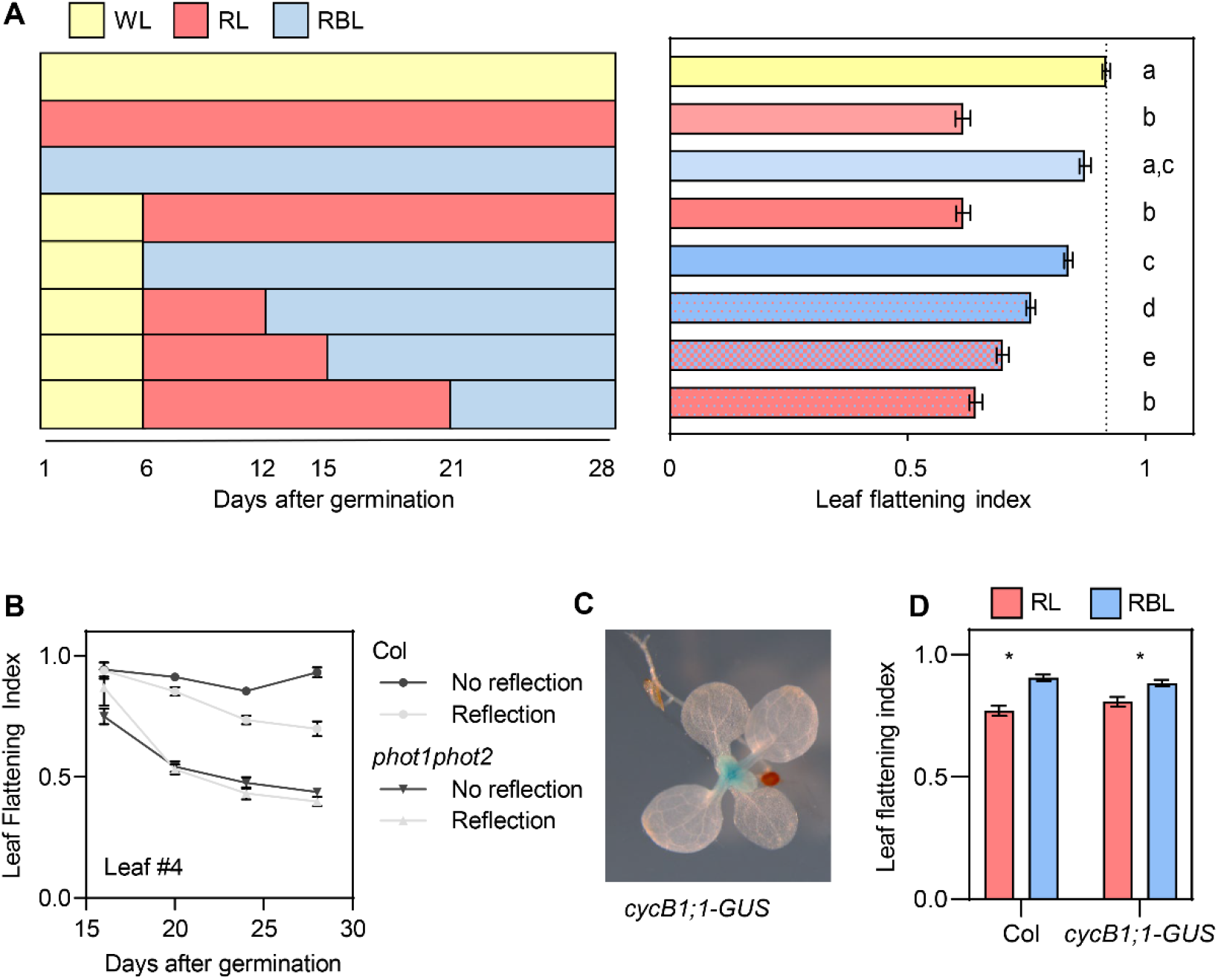
Phototropins control leaf shape reversibly during the expansion phase. A- Leaf flattening index of Col plants grown in WL on soil and transferred to WL, RL or RBL at different time points. Measurements were done 28 days after germination. Bars represent mean ±SE of 40 – 48 leaves. Different letters represent significant differences among means (p<0.05) in an ANOVA followed by Tukey’s test. Left: scheme of the treatments. B- Leaf flattening index in Col and phot1phot2 plants grown on soil covered with dark (No reflection) or clear (Reflection) aluminum. Results correspond to leaf 4 during development since leaves are big enough to be measured. Each point represents mean ±SE 8 to 11 plants. C- GUS staining of plants expressing *cycB1;1-GUS* grown on soil covered with clear (Reflection) or dark (No reflection) aluminum foil. Staining was performed 2 weeks after germination, when leaf 1 and 2 were more than 3mm long, but before they respond to the light stimulus. D- Leaf flattening index in plants expressing *cycB1;1-GUS* grown in WL for two weeks and transferred to RL or RBL for one week. Bars represent mean ±SE of 16 - 22 leaves. * p < 0.05 in ANOVA followed by Dunnett’s multiple comparison test.

### Phototropins regulate the auxin signaling pattern in the medio-lateral and abaxial-adaxial axes of the leaf blade

In the case of stem phototropism, differential cell expansion that causes bending is regulated by auxin, with higher auxin levels on the shaded than on the lit side of the hypocotyl (Ding et al., 2011; Willige et al., 2013; Legris and Boccaccini, 2020). To evaluate whether in the leaf phototropins also regulate auxin signaling, we determined the distribution pattern of the auxin output reporter *pDR5rev::3XVENUS-N7* (*DR5:VENUS*) (Heisler et al., 2005) in the blade of expanding leaves using epifluorescence microscopy. In Col plants grown in WL the DR5:VENUS signal was higher on the abaxial side of the blade than on the adaxial side, where it was restricted to the margins (Fig. 5). We observed a correlation between the *DR5* expression pattern and light-regulated flattening. In conditions where leaves curled downwards, i.e. reflected light from below or the absence of BL (RL), the VENUS signal increased on the adaxial side (Fig. 5A, B). Adding low intensities of BL, which was enough to promote flattening (Fig. 1), reduced the VENUS signal on the adaxial side of the blade and restored the distribution pattern found in WL (Fig. 5A). This response depended on phototropins, since the double *phot1phot2* mutant had an altered *DR5:VENUS* expression pattern in WL and did not respond to the light treatments (Fig. 5C, D). These results are consistent with a model where phototropins create an auxin response gradient in the leaf. However, in transverse cuts of fixed leaves expressing *DR5:VENUS* we only detected VENUS signal in the epidermis. In plants grown in WL no signal was detected in the adaxial epidermis in the middle of the blade, while in those grown in RL we detected it in both epidermal layers (Fig. S5). Taken together with our data indicating that *pML1:PHOT1* complemented *phot1phot2* (Fig. 2) these results suggest a particularly important role of phototropin activation and auxin signaling on both epidermal layers to control leaf flattening. In particular, BL-activated phototropins in the adaxial epidermis inhibit auxin signaling in the center of the blade to promote flattening.

**Figure 5 –.**
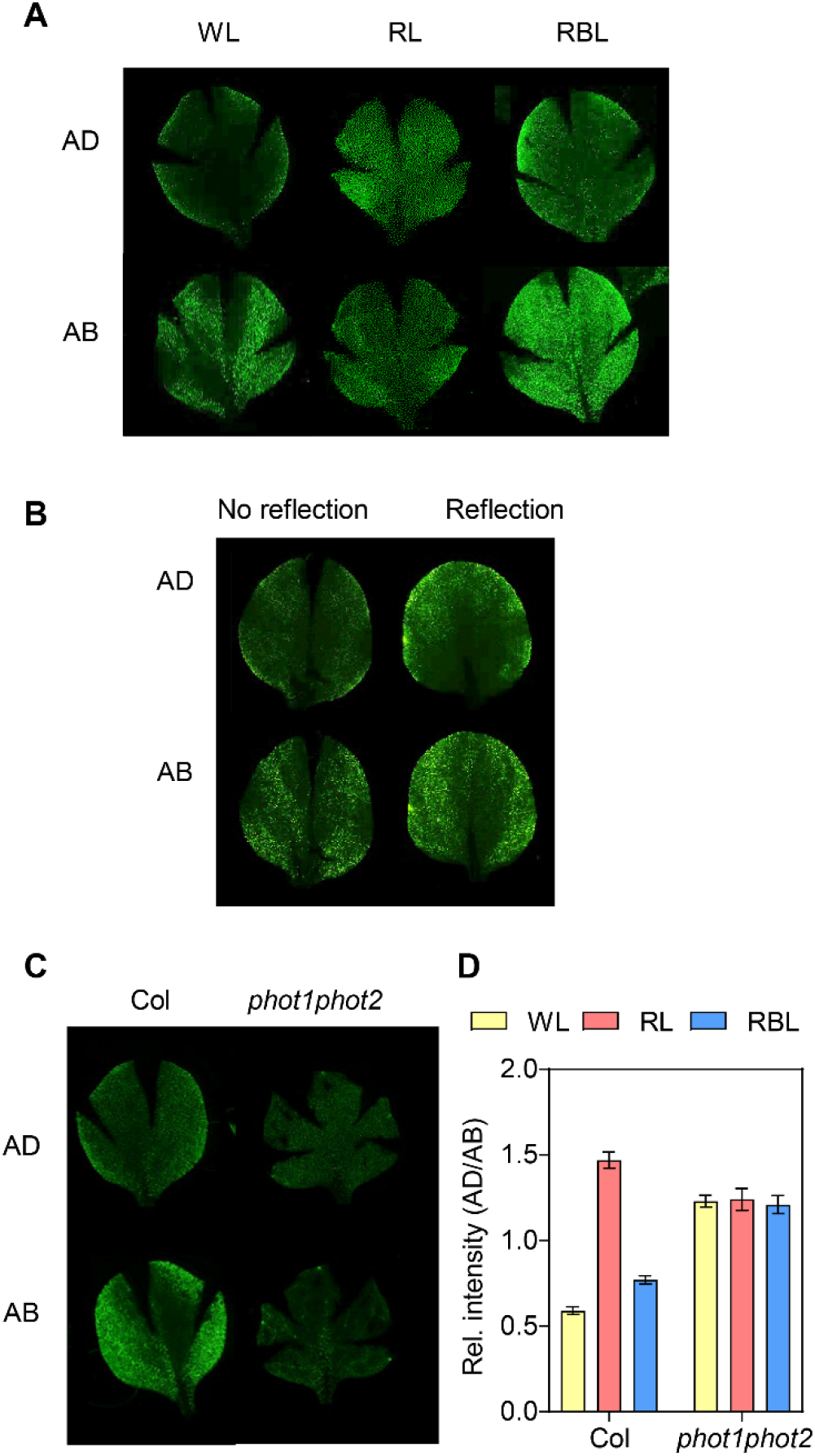
Leaf shape correlates with the auxin signaling pattern in the blade. A- Epifluorescence images of leaf blades of Col plants expressing DR5:VENUS grown in WL for two weeks and transferred for one week to WL, RL, RBL. For each leaf images were taken on the adaxial (AD) and abaxial (AB) side. B- Epifluorescence images of leaf blades of Col plants expressing DR5:VENUS grown in WL on soil covered with dark (No reflection) or clear (Reflection) aluminum foil for three weeks. For each leaf images were taken on the AD and AB side. C- Epifluorescence images of leaf blades of Col and phot1phot2 plants expressing DR5:VENUS grown in WL for two weeks and transferred for one week to WL, RL, RBL. For each leaf images were taken on the AD and AB side. D- Quantification the VENUS fluorescence in epifluorescence images of whole blades. For each leaf images were taken on the AD and AB side, DR5:VENUS fluorescence was quantified and the ratio between fluorescence in each side was calculated. Each bar represents mean ±SE of 10 to 16 leaves.

### Light-regulated leaf flattening involves auxin transport

Changes in the *DR5* promoter activity could be due to changes in auxin synthesis, transport or sensitivity. Given that auxin transport has a strong role in hypocotyl phototropism (Christie et al., 2011; Ding et al., 2011; Willige et al., 2013), we tested whether this was also the case during leaf flattening. First, we used the auxin transport inhibitor N-1-naphthylphthalamic acid (NPA) to block auxin transport during the leaf expansion phase. Treating plants with this compound in WL-grown plants had a similar effect on leaf flattening as inactivating phototropins with a RL treatment (Fig. 6A). This phenotypic response correlated with changes in the *DR5:VENUS* pattern: treatment with NPA increasing the VENUS signal in the leaf (Fig. 6B). To determine which transporters play a role in this response, we grew mutants affected in the major classes of auxin transporters. All tested auxin transporters families played a role in leaf flattening when plants were grown in standard conditions (Fig. 6C, D, E). To evaluate whether this defect was related to light-regulated leaf development, we measured LFI in response to light signals in these mutants. *abcb1 abcb19* mutants responded to light irradiated from the top or from below (Fig. 6D, E), indicating that these auxin transporters were not essential to promote leaf flattening by BL. However, they showed a regulatory role in the response, since this double mutant responded more to light from below as shown by the significant interaction term in the ANOVA (Fig. 6E). In agreement with the observation that NPA inhibits BL-regulated leaf flattening, *pin3pin4pin7* triple mutants did not respond to light from the top and showed a reduced response to reflected light (Fig. 6D, E). Interestingly, among the different auxin transport mutants tested, *aux1lax1* was the only one showing a striking resemblance to the *phot1phot2* rosette morphology (Fig. 6C). Also, this mutant showed a very similar flattening defect as *phot1phot2*, with no response to BL from above, and a very reduced response to reflected light (Fig. 6D, E). Overall, these results are consistent with a role of phototropins regulating auxin transport to control leaf flattening and uncover a potentially novel mechanism of phototropin signaling involving AUX1/LAX auxin transporters.

**Figure 6 –.**
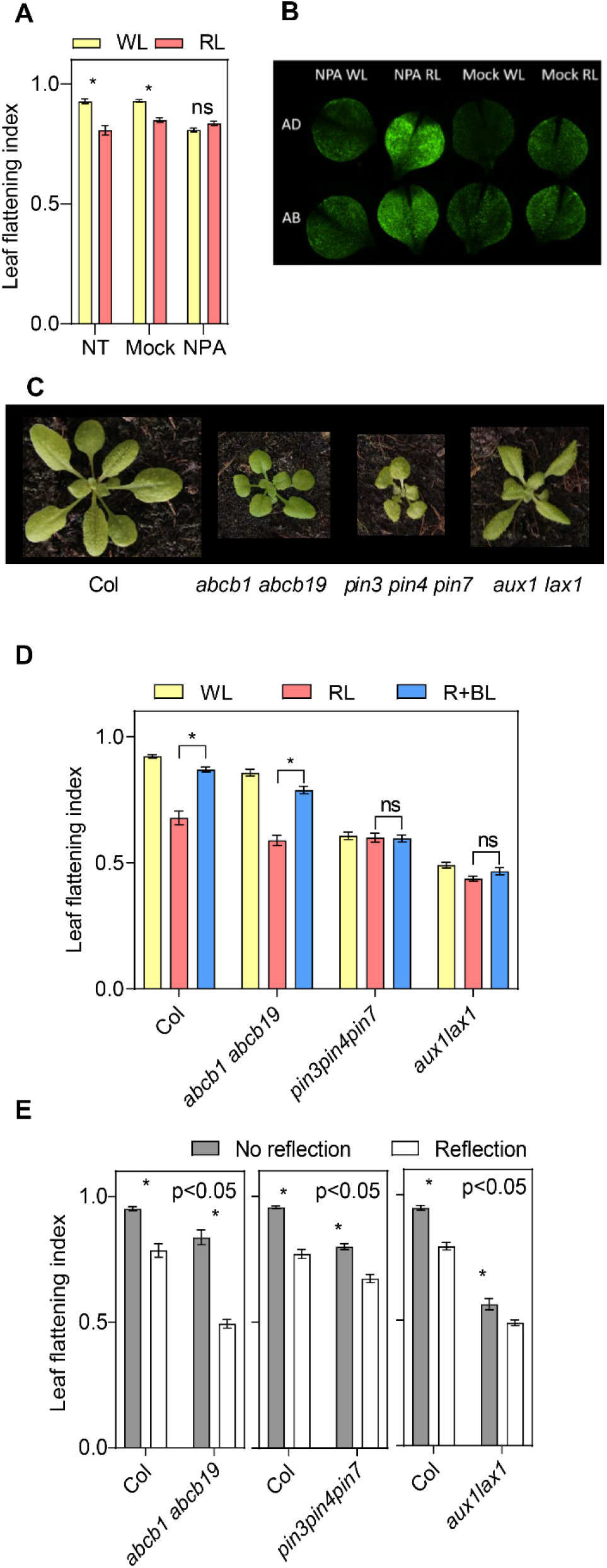
Auxin transporters are required for light-regulated leaf flattening. A- Leaf flattening index of Col plants grown in WL for two weeks and transferred to WL or RL for one week. During this week some plants were sprayed with NPA, mock solution (Mock) or not treated (NT). Bars represent mean ±SE of 30 – 80 plants. B- Epifluorescence images of leaf blades of Col plants expressing DR5:VENUS grown in WL for two weeks and transferred for one week to WL or RL for one week while being treated with NPA or mock. For each leaf images were taken on the adaxial (AD) and abaxial (AB) side. C- Pictures of Col and various auxin transporters mutants grown in WL for three weeks. D- Leaf flattening index in plants of various genotypes grown on soil for two weeks and transferred to WL, RL or RBL for one week. Bars represent mean ±SE of 30 – 54 plants. * p < 0.05 in ANOVA followed by Dunnett’s multiple comparison test. F- Leaf flattening index in plants of various genotypes grown in WL on soil covered with dark (No reflection) or clear (Reflection) aluminum foil for three weeks. Bars represent mean ±SE of 14 - 44 plants. *p < 0.05 in ANOVA followed by Dunnett’s multiple comparison test. Significance of the interaction term (Genotype x Treatment) in the ANOVA is shown in the top right corner in each case.

## DISCUSSION

### Light direction perceived by phototropins is a key aspect regulating leaf blade flattening

Leaves are the main light harvesting organs in plants, and their shape directly influences photosynthesis through light interception and gas exchange. Leaf flattening is strongly controlled by the developmental program of the plant, from early patterning through expansion (Chitwood and Sinha, 2016; Xiong and Jiao, 2019; Heisler and Byrne, 2020). Given the tight relationship between leaf shape and light interception, we focused on how BL signals control leaf flattening, and found that phototropins perceive light direction to control leaf shape accordingly (Fig. 1, Fig.2). This effect, combined with the role of phototropins on leaf positioning and anatomy, indicate strong interactions between BL signaling and endogenous cues in controlling leaf development according to the light environment (Inoue et al., 2008; de Carbonnel et al., 2010; Kozuka et al., 2011). Indeed, phototropin mutants show reduced biomass accumulation, which can at least in part be explained by their leaf morphology (Takemiya et al., 2005; Inoue et al., 2008; de Carbonnel et al., 2010). Interestingly, the response to light direction in the leaf is conserved in lettuce, where leaves irradiated on the abaxial side showed a curled phenotype which also correlated with reduced biomass (Wang et al., 2021).

Phototropins control leaf flattening reversibly, late in development, when cell division has ended, but while leaves are still expanding (Fig. 4). This role of light on blade expansion presumably occurs in the epidermis, as this tissue has major effects on growth regulation (Savaldi-Goldstein et al., 2007). This is consistent with complementation of *phot1phot2* leaf curling by epidermal expression of PHOT1-citrine (Fig. 2). Moreover, in lettuce light from below affects cell shape and expansion in the adaxial and abaxial epidermis to regulate leaf curvature (Wang et al., 2021). phot2 regulates palisade mesophyll development in a light-dependent manner (Kozuka et al., 2011). However, we observed changes in leaf flattening in response to low BL and in *phot2* mutants (Fig. 1). In addition, plants expressing phot2 in the epidermis respond to light to control leaf flattening but not to regulate palisade mesophyll development (Kozuka et al., 2011). We conclude that phototropins play a primary role in the epidermis to control blade flattening.

Leaf flattening correlates with the pattern of auxin signaling reporter *DR5:VENUS*. In conditions where leaves were flat, the *DR5:VENUS* signal was high in the margins and in the abaxial epidermis, while downwards curling correlated with an increase in the *DR5:VENUS* signal in the center of the adaxial epidermis (Fig. 5). This pattern depended on BL and phototropins (Fig 5D). We cannot exclude the possibility that an auxin pattern in the adaxial epidermis (margins vs. center) also explains leaf flattening, since both aspects were affected in *phot1phot2*. The role of abaxial auxin concentration on leaf flattening has been studied in earlier stages of leaf development (Guan et al., 2017; Xiong and Jiao, 2019; Heisler and Byrne, 2020). Although some controversy still exists on whether higher auxin in the abaxial side of the primordium is an early signal creating the adaxial-abaxial patterning, it is possible that this signal helps maintain tissue patterning (Heisler and Byrne, 2020). It would be interesting to know whether phototropins affect this auxin distribution early in development. However, we note that the *phot1phot2* leaf flattening phenotype is most obvious late in development highlighting their importance late during leaf expansion (Fig. 4B). Whether changes in the *DR5* expression pattern reflect changes in auxin levels or downstream signaling requires further investigations. NPH4/ARF7, which is required for hypocotyl phototropism, is developmentally regulated showing expression predominantly in the adaxial side of the primordia, as most activator ARFs (Guan et al., 2017). Moreover, *arf7* mutants show deficient leaf flattening, with a phenotype similar to *phot1phot2* suggesting a role for auxin signaling in light-controlled leaf flattening (Watahiki and Yamamoto, 1997).

All auxin transporter mutants tested showed defects in leaf flattening and in response to light (Fig. 6). However, the only mutant resembling the *phot1phot2* phenotype was *aux1lax1* (Fig. 6). AUX/LAX proteins control phyllotaxis, venation and margin development (Reinhardt et al., 2003; Swarup and Bhosale, 2019; Xiong and Jiao, 2019). Interestingly, these genes also control leaf flattening in tomato (Pulungan et al., 2018). PINs have many roles in leaf development, including primordia establishment, phyllotaxis and early patterning (Xiong and Jiao, 2019; Heisler and Byrne, 2020). The triple *pin3pin4pin7* mutant did not respond to light signals, suggesting that these auxin exporters are essential to regulate leaf flattening (Fig. 6). Nevertheless, the leaf and rosette morphology of this mutant differs strongly from *phot1phot2* (Fig. 6C). Given the many roles of PIN auxin transporters in development, it is possible that earlier defects in leaf patterning render the mutants unable to respond to light signals. However, inhibiting PINs with NPA later in development caused similar flattening defects as inactivating phototropins indicating that during blade expansion polar auxin transport is required for leaf flattening (Fig. 6A). ABCB1 and ABCB19 are involved in phototropin-dependent leaf positioning and flattening (Jenness et al., 2020). In our experiments, and consistent with previous reports, the *abcb1abcb19* mutant had a flattening defect in WL (Fig. 6). However, this mutant responded to light signals from the top (Fig. 6D) and it responded to light from below more than the wild type (Fig. 6E). Hence, in the conditions analyzed here, ABCB1 and ABCB19 are not required for phototropin-mediated leaf flattening, but they have a regulatory role. We conclude that multiple auxin transporters regulate leaf flattening but AUX/LAX play a particularly important role. This contrasts with the minor role of AUX/LAX in hypocotyl phototropism (Christie et al., 2011; Hohm et al., 2014), suggesting that somewhat different mechanisms control phot-mediated auxin gradients formation in hypocotyls versus leaves.

### Conserved aspects and differences of phototropin signaling between leaf blades and hypocotyls

Our study reveals that general principles of phototropin action are conserved between stem phototropism and leaf flattening, in particular when light reaches the adaxial side of the leaf. In both cases phototropins have the capacity to perceive light direction, and to regulate auxin signaling to change cell expansion and ultimately organ curvature (Fig. 1, Fig 4, Fig 5) (Legris and Boccaccini, 2020). In hypocotyl phototropism phot1 has a major role in response to low BL intensities (less than 1μmol.m^−2^.s^−1^) and phot2 responds to higher light intensities. Our results in leaves indicate that both phot1 and phot2 respond to BL intensities below 0.4μmol. m^−2^. s^−1^ in the adaxial and in the abaxial side, although phot1 has a more prominent role in response to low light intensities coming from the top (Fig. 1). The current model for hypocotyl phototropism is that light coming from a predominant direction establishes a light gradient across the organ leading to a phototropin activation gradient, which determines bending orientation. The situation in leaves is different because the abaxial and adaxial sides respond differently to BL (Fig. 1). The lower side is more sensitive to light than the upper side, since low BL applied to the abaxial side triggers bending despite high BL reaching the adaxial side (Fig. 1B). phot1-mediated sensing on both sides of the leaf was further established using *PHOT1-citrine* under the control of *pAS2* and *pFIL* promoters (Fig. 2). Moreover, we could determine that in response to BL phot1 is downregulated (Fig. 3), which is linked to desensitization in hypocotyls (Roberts et al., 2011). The higher light exposure of the adaxial side may trigger asymmetric phot1 degradation thereby explaining the higher light sensitivity of the lower side of the leaf.

Differential sensitivity to light in the adaxial and abaxial sides of the leaf may also be caused by differential signaling mechanisms on these developmentally distinct parts of the leaf. In line with this idea, the *nph3* mutant has distinct phenotypes depending on the side of leaf irradiation (Fig.3). Consistent with previous data, NPH3 is required to promote leaf flattening in response to LB irradiation from the top (Fig. 3)(Inoue et al., 2008). In contrast, NPH3 may inhibit the BL response on the abaxial side (Fig. 3D, E). NPH3 is an essential positive element of phototropin signaling during hypocotyl phototropism (Christie et al., 2018). It is therefore surprising that NPH3 may counteract phototropin signaling depending on the tissue type. One alternative interpretation of the *nph3* leaf flattening phenotype is that blade curvature is a balance between light responses on both sides of the leaf. In the absence of NPH3 phototropin signaling in the adaxial side of the leaf could be impaired, but response to light from below could be normal, causing an imbalance and explaining the *nph3* phenotype. Irrespective of the mechanism, our results show that phototropins can perceive and respond to light in the abaxial epidermis in an NPH3-independent manner possibly involving other members of the NRL family such as RPT2, which also regulates leaf flattening (Harada et al., 2012).

In the hypocotyl PKS1, PKS2 and PKS4 have a major role regulating phototropism (Kami et al., 2014). Here we found that PKS3 is the main PKS protein involved in leaf flattening (Fig. 3). Single *pks3* mutants showed a flattening defect comparable to *phot1phot2* and reduced responses to light signals. However, as observed for hypocotyl phototropism, there is functional redundancy among the members of the PKS family to control BL responses of leaves (Fig. 5). This is consistent with the reported roles in leaf flattening and positioning of PKS1 and PKS2, and the flattening defect found in *pks1pks2pks4* triple mutant (de Carbonnel et al., 2010). PKS2 functions in phot2 signaling controlling flattening (de Carbonnel et al., 2010). In our experiments *pks3* mutants had a reduced but significant response to low blue light, suggesting that the phot1 pathway was still partially active in this mutant (Fig. 3). Interestingly, PKS proteins were previously shown to act at the interface between internal and external cues. During hypocotyl phototropism in etiolated seedlings the response of *pks* mutants depends on whether light reaches the cotyledons or the hook (Kami et al., 2014). The results presented here for PKS3 in leaf flattening further show the central role of PKS proteins in coordinating plant development with the environment.

## MATERIALS AND METHODS

### Plant material

The following Arabidopsis (Arabidopsis thaliana; Col-0) mutants were used previously: *phot1-5*, *phot2-1*, *phot1-5 phot2-1*, *nph3-6* (de Carbonnel et al., 2010); *phyB-9*, *phyB-9 ven4-bnen* (Yoshida et al., 2018), *cry1-304*, *cry2-1*, *cry1-304 cry2-1* (Boccaccini et al., 2020); *aux1-21 lax2-1* (Hohm et al., 2014), *pin3-3 pin4-101 pin7-101* (Willige et al., 2013), *pgp19-101* (Jenness et al., 2019).

The following transgenic lines were described before: *pML1:PHOT1-citrine*, *pPHOT1:PHOT1-citrine* (Preuten et al., 2013), *pCER6:PHOT2-GFP* (Kozuka et al., 2011), *pCYCB1;1::NterCYCB1-GUS* (Marrocco et al., 2009), *pDR5rev::3XVENUS-N7* (Heisler et al., 2005).

*pDR5rev::3XVENUS-N7* in *phot1-5 phot2-1*, as well as multiple *pks* mutants containing *pks3-9*, were obtained by crossing. The methods used to genotype the mutations appear in Supplemental table 1 and Supplemental table 2. The presence of the *pDR5rev::3XVENUS-N7* transgene was selected by resistance to BASTA.

### Generation of CRISPR mutant alleles of *PKS3*

To create *pks3-9* and *pks3-10* mutants the protocol by Hyun et al (Hyun et al., 2015) was followed. pYB196 and pRG-ext-CCR5 were provided by George Coupland. The sequence of the guide used was 5’-AGATCATGTTGATTCCACGG-3’.

The selected mutant alleles were named *pks3-9* and *pks3-10*. *pks3-9* has a 2-bp deletion and a mutation in position 192 (Col sequence 5’-TGATTCCA-3’, mutant allele 5’-CGTTTT-3’) which creates an early stop codon resulting in a 73-aminoacid protein. *pks3-10* has a base insertion (A) in position 199, which creates an early stop codon and a truncated 74-aminoacid protein.

### Generation of transgenic lines expressing PHOT1 under*pAS2* and*pFIL* promoters

For pAS2::PHOT1-citrine the backbone and CDS PHOT1-citrine were obtained from pPHOT1::PHOT1-citrine (Preuten et al., 2013). The pAS promoter was amplified from pCRII-TOPO pAS2 (Wu et al., 2008). For pFIL::PHOT1-citrine the backbone and pFIL promoter were obtained from pGWB-NB1-pFIL (Tameshige et al., 2013). The PHOT1-citrine CDS was obtained from pPHOT1::PHOT1-citrine (Preuten et al., 2013). Transgenic plants were generated by introduction of the plant expression constructs into pSOUP-containing *Agrobacterium tumefaciens* strain GV3101. Transformation of *phot1-5phot2-1* plants was done by floral dipping. Based on segregation of Basta-resistance, homozygous T3 lines with a single transgene were selected.

### RT-qPCR

Plants were grown on plates for 12 days in continuous light. For RNA extraction only the aerial part containing hypocotyls, the shoot apical meristem and small leaves of ~ 1mm long were grinded in liquid nitrogen. RNA extraction was performed with the RNAeasy plant mini kit (Quiagen). Each sample consists of 6 plants, except for pFIL-9 (4 plants) and pFIL-37 (8 plants), and three replicates per line were harvested. The reverse transcription was done using SuperScript II (Invitrogen) from 1μg of RNA. Transgene expression levels were measured quantifying the citrine transcript with primers LAP038 and LAP045. Three housekeeping genes were used: GAPDH, UBC, YLS8. Primers used for this analysis are listed in Supplemental table 2.

### Growth conditions

Seeds were sowed on soil (CL TON KOKOS, CLASSIC Profisubstrat, Einheits erde) in square plastic pots (8 cm x 8 cm × 7 cm, width × depth × height) watered with Solbac solution 1:400 (Andemat Biocontrol). After stratifying for 3-5 days plants were grown in 16:8 light/dark cycles in a walk-in incubator equipped with white and red LEDs, PAR =100 μmol.m^−2^.s^−1^, at 22°C.

For experiments involving light reflection seeds were sowed on 0,8% water-agar plates, stratified for 3-5 days and grown for 4 days in light until they were de-etiolated. A square of aluminum foil was placed over the soil, and four small holes (2-3 mm diameter) were done on each corner. The holes were filled with moistured soil, and seedlings were transplanted inside.

In Fig. 1A, B, D seeds were surface-sterilized and sowed in square transparent plates (10cm × 10cm × 2cm) with half MS media (Duchefa) plus 0,8% agar (Roth).

### Light treatments

White light (WL), red light (RL) and red + blue light (RBL) treatments were performed in a Percival incubator, model AR36-L3, set to 16:8 light/dark cycles, 22°C, relative humidity 70%. WL was obtained with fluorescent tubes (OSRAM Lumilux cool white L18W/840), RL was obtained with red LEDs and RB was obtained with red LEDs combined with fluorescent tubes. PAR was set to 100 μmol.m^−2^.s^−1^ in all cases. The red, blue and far-red values (μmol.m^−2^.s^−1^) were: WL = 20; 7; 1,45; RL= 60; 0,003; 0,1; RBL = 60; 0,4; 0,1.

In Fig. 1A, 1B, 1D plants were grown on plates in Percival incubators model AR22-LX, set to 16:8 light/dark cycles, 22°C. WL was obtained with fluorescent tubes (OSRAM Lumilux cool white L18W/840), RL was obtained with red LEDs. Light from below was obtained with LEDs (FloraLED, CLF plant climatics).

Light intensities were measured with a radiometer IL 1400A (International Light, US) using a W filter (#9540) and PAR (#21777), TBLU (#21853), TRED (#22237) and TFR (#22238) filters.

### Leaf flattening index measurement

Leaf blades were placed on top of an agar plate with the adaxial face up and imaged. Then they were flattened performing cuts in the margins and sticking the adaxial side on transparent scotch tape. The flattened leaves were imaged again. Image analysis was performed with Image J. Pictures were converted to 8-bit and a manual threshold was applied to segment the whole blade. The whole blade was selected using the Wand (tracing) tool. Leaf flattening index was calculated as the ratio Area (before flattening) / Area (after flattening) for each leaf. Unless stated otherwise, all measurements were done on leaf 1 and 2.

### Fluorescence microscopy

Confocal microscopy images were taken with an LSM 710 confocal microscope (Zeiss). For imaging DR5:VENUS, pPHOT1:PHOT1-citrine, pAS2:PHOT1-citrine and pFIL:PHOT1-citrine samples were excited with an Argon laser (514nm) and detection was done between 520 and 560nm. For pCER6:PHOT2-GFP samples were excited with an Argon laser (488nm) and detection was done between 505nm and 530nm. For the calcofluor white stain samples were excited with a 405nm laser and detection was done between 420 and 470nm. In all cases a channel was set to detect chlorophyll, exciting with the laser used for excitation of the fluorophore of interest, and detecting between 600 and 750nm.

For Fig. S5 leaves were fixed in 4% PFA for two hours, washed with 1X PBS and stained with calcofluor white (0,1%) in PBS applying vacuum and staining overnight at room temperature. Stained leaves were embedded in 1,5% agarose, cut with a razor blade and mounted on a glass bottom dish with PBS.

For Fig. 5 and Fig. 6A, leaves were harvested and immediately mounted between two coverslips with water, doing cuts in the margins to allow flattening. For each leaf images were taken on the adaxial and abaxial side using a Leica M205 FCA stereomicroscope equipped with a GFP filterset (excitation 470/40, emission 525/50).

For Fig. 5D DR5:VENUS fluorescence from whole blades was quantified using ImageJ, applying a threshold to segment the whole blade and selecting it with the wand (tracing) tool.

### GUS staining

Leaves were harvested and fixed in 90% acetone for 4h and rinsed twice with 50Mm NaPO4 before vacuum infiltrating with the staining solution (4 times 15min each). The staining solution contained: EDTA pH 8,5 10mM, NaHPO4 50mM, Triton C-100 0,1%, X-Gluc 0,5 mg/ml in water. Samples were incubated at 37°C overnight and cleared with 70% EtOH at 4°C, over various days changing EtOH regularly. Images were taken with a Leica M205 FCA stereomicroscope.

### Western blot

Plants were grown for two weeks and transferred to WL, RL or RBL at CT8 for 24h. Leaf blades were harvested on liquid nitrogen, each sample consisting of leaves 1 and 2 from one plant.

Total proteins were extracted by grinding the seedlings in 50μl 2x Laemmli buffer with 10% β-mercaptoethanol. Samples were heated 5min at 95°C and centrifuged for 1min. 10μl per sample were loaded in a pre-cast 4-15% gradient agarose gel (Mini-PROTEAN TGX, BioRad). Electrophoresis was performed at 100mV during 75min. Transfer was performed using the TransBlot Turbo system (BioRad). Detection of PHOT1-citrine was done using Living Colors anti-GFP antibody JL-8 (632381, Clontech), 1:4000 in PBS 5% milk 0,1% tween (PBSTM). Detection of endogenous PHOT1 was done using anti-PHOT1 1:5000 in PBSTM (Christie et al., 1998; Cho et al., 2007). DET3 was used as a loading control, and detected using anti-DET3 1:20000 in PBSTM (Schumacher et al., 1999), as well as Histone 3 which was detected using anti-Histone H3 (ab1791, abcam) 1:3000 in PBSTM. Membranes were blocked for 1h at room temperature with PBSTM and incubated overnight with the corresponding primary antibody at 4°C. anti-PHOT1, anti-DET3 and anti-H3 were detected with anti-rabbit IgG, HRP conjugate (W4011, Promega) 1:2500 in PBSTM. anti-GFP was detected with anti-mouse IgG, HRP conjugate (W4021, Promega) 1:5000 in PBSTM. Membranes were revealed using Immobilion Western Chemiluminiscent HRP substrate (WBKLS0500, Millipore). Chemiluminiscence was detected using an ImageQuant LAS 4000 mini (GE Healthcare). Image quantification was performed using ImageJ.

### NPA treatment

A stock solution of Naphthylphtahalamic acid (NPA, Prod No. N0926.0250, Duchefa) was prepared to a concentration of 10 mM in DMSO and kept at −20°C. The day of the treatment a new dilution was performed in water plus 0.15% Tween20 to a final concentration of 10μM. The mock solution consisted of 1:1000 DMSO in 0.15% Tween20. NPA was sprayed from above the day plants were transferred to RL. Leaf flattening index was measured after 7 days.

## Acknowledgements

We thank Prof. George Coupland for sharing the plasmids used for CRISPR, Prof. John Christie for the anti-PHOT1 antibody, and Prof. Akira Nagatani for sharing seeds carrying *pCER6:PHOT2-GFP*. The plasmid pCRII-TOPO pAS2 was provided by Prof. Patricia Springer, and the plasmid pGWB-NB1-pFIL by Prof. Kiyoshi Tatematsu. Figure 6A was done with the collaboration of the master student Laetitia Tcheng. We thank the CIF facility at the University of Lausanne for assistance with confocal microscopy.

## Accession Numbers

The Arabidopsis Genome Initiative numbers for the genes mentioned in this article are as follows: AT2G18790 (PHYB), AT3G45780 (PHOT1), AT5G58140 (PHOT2), AT4G08920 (CRY1), AT1G04400 (CRY2), AT2G02950 (PKS1), AT1G14280 (PKS2), AT1G18810 (PKS3), AT5G04190 (PKS4), AT5G64330 (NPH3), AT2G36910 (ABCB1), AT3G28860 (ABCB19), AT1G70940 (PIN3), AT2G01420 (PIN4), AT1G23080 (PIN7), AT2G38120 (AUX1), AT5G01240 (LAX1), AT4G21750 (ML1), AT1G68530 (CER6), AT1G65620 (AS2), AT1G68530 (FIL), AT1G13440 (GAPDH), AT5G25760 (UBC), AT5G08290 (YLS8).

---

**Supplemental figure S1** – Light treatments and growth conditions

A- Dark aluminum foil does not affect leaf shape compared to dark soil. Col plants grown for three weeks in WL in soil or in soil covered with dark aluminum foil. Leaf flattening index (mean ±SE) of 22-32 leaves.

B- Images of Col plants grown in various conditions. Note that the presence of dark aluminum foil does not affect plant morphology, while plants grown in plates have smaller round leaf blades, with irregular surfaces, and thinner petioles than those grown in soil.

C- Spectra of reflected light by colored aluminum foil. Colored aluminum foil was placed in front of a light source and the reflected light was measured. Top left, spectrum of the light source. To facilitate comparison of the shape of the spectra among colors in each case the spectrum plotted is relative to the maximum value.

D- The *ven4-bnen* mutation does not affect phyB leaf flattening. Plants grown for two weeks in WL and transferred to WL, RL or RBL for one week. Left panel: *phyB-9 ven4-bnen*. Right panel: *phyB-9* without the *ven4* mutation. Bars represent mean ±SE of 22-28 leaves (left) and 37-48 leaves (right).

**Supplemental figure S2** – Expression levels and distribution pattern of pAS2 and pFIL lines

A- Expression levels of *pAS2* and *pFIL* lines determined by qPCR (waiting for details from Laure). Each bar represents mean ±SE of three biological replicates.

B- Confocal microscopy images of leaf blades of plants expressing *PHOT1-citrine* under various promoters. The same leaf was imaged on the adaxial and the abaxial side.

**Supplemental figure S3** – PHOT1 levels decrease with BL

A- PHOT1 levels decrease with blue light in Col Plants grown for two weeks in WL and transferred to WL, RL or RBL for one day. Membranes were probed with anti-PHOT1 and anti-DET3 or anti-Histone3 (H3) antibodies. Top: Experiment quantified in Fig. 3A. Bottom: Independent experiment and its quantification. Each bar represents mean ±SE of four replicates.

B- Plants expressing *pPHOT1:PHOT1-citrine* grown for two weeks in WL and transferred to WL, RL or RBL for one day. Membranes were probed with anti-GFP and anti-DET3 antibodies. Left: Quantification of the western blot. Each bar represents mean ±SE of four replicates.

**Supplemental figure S4** – Phototropins control flattening reversibly and late in development

Leaf flattening index of Col plants grown on soil and transferred to WL, RL or RBL at different time points. Measurements were done 28 days after germination. Bars represent mean ±SE of 24 – 48 leaves. Left: scheme of the treatments.

**Supplemental figure S5** - DR5:VENUS signal is detected in the epidermis.

Col plants expressing nuclear-localized VENUS under the control of the DR5 promoter (green) were grown for two weeks in WL and transferred to WL or RL for one week. Leaves 1 and 2 were fixed and stained with calcofluor white (grays). Adaxial and abaxial epidermal layers are labeled as AD and AB respectively.

**Supplemental table S1 -** Genotyping conditions to select mutations in crosses

**Supplemental table S2 –** Primers used in this study

